# Bacterial heterozygosity promotes survival under multidrug selection

**DOI:** 10.1101/2024.05.09.593358

**Authors:** Shraddha Shitut, Thomas van Dijk, Dennis Claessen, Daniel Rozen

## Abstract

Although bacterial cells typically contain a single chromosome, some species are naturally polyploid and carry multiple copies of their chromosome. Polyploid chromosomes can be identical or heterogeneous, the latter giving rise to bacterial heterozygosity. While the benefits of heterozygosity are well studied in eukaryotes, its consequences in bacteria are unknown. Here we examine this question in the context of antibiotic resistance to understand how bacterial heterozygosity affects bacterial survival. Using a cell wall deficient model system in the actinomycete *Kitasatospora viridifaciens*, we found that heterozygous cells containing different chromosomes expressing different antibiotic resistance markers persist across a broader range of antibiotic concentrations. Recombinant cells containing the same resistance genes on a single chromosome also survive these conditions, but these cells pay a significant fitness cost due to constitutive expression of these genes. By contrast, heterozygous cells mitigate these costs by flexibly adjusting the ratio of their different chromosomes, thereby allowing rapid responses in temporally and spatially variable environments. Our results provide evidence that bacterial heterozygosity can increase adaptive plasticity in bacterial cells, in a similar manner to the evolutionary benefits provided by multicopy plasmids in bacteria.

**Graphical abstract:** 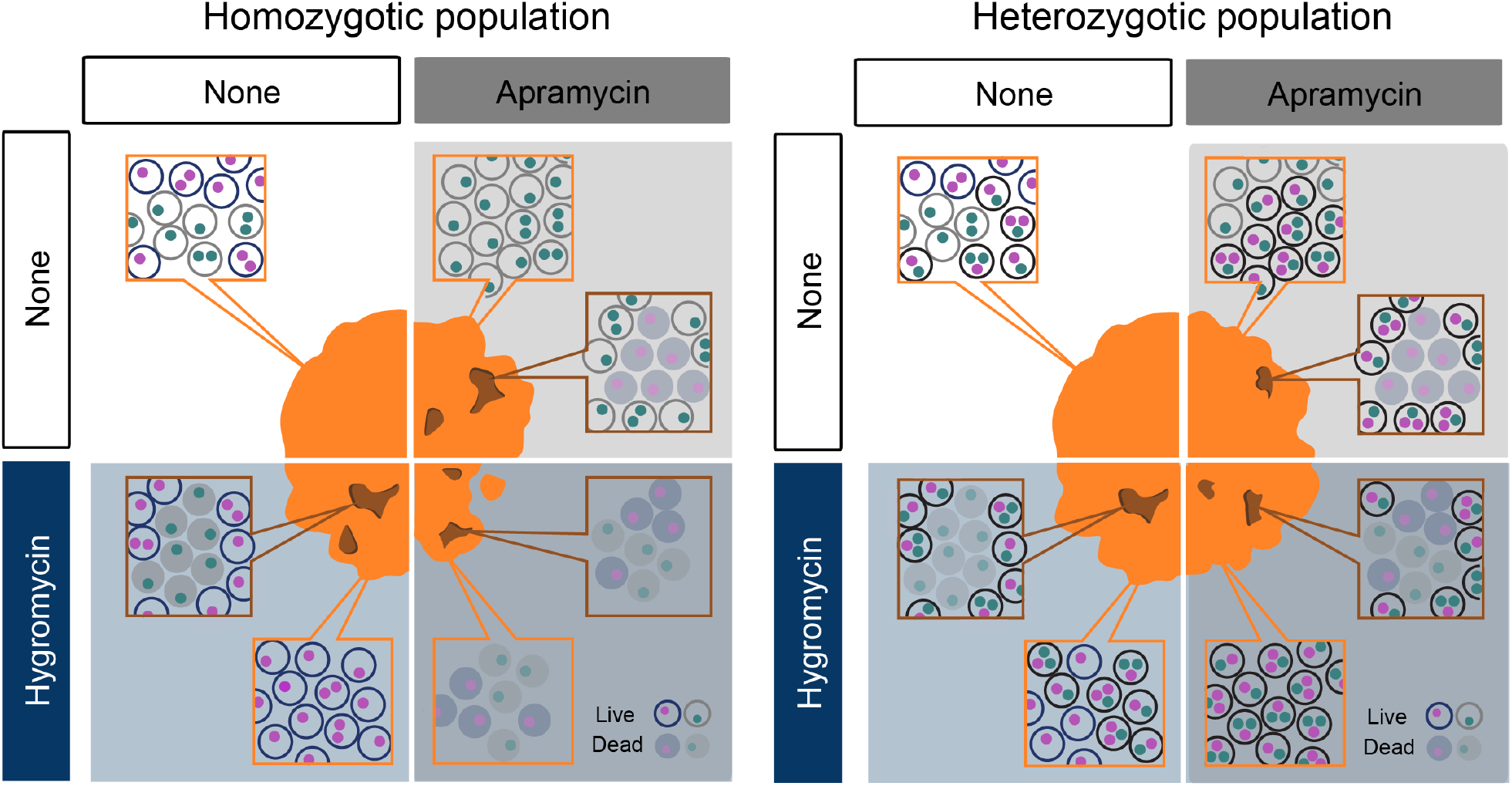

- Heterozygotic bacterial cells (polyploid with multiple chromosome types) can persist across a wider range of conditions than homozygotic cells.
- Heterozygous cells can adaptively adjust chromosomal frequencies in response to external conditions
- Flexibly adjusting chromosome ratios enables heterozygous cells to mitigate the costs of antibiotic resistance and allows rapid responses to temporally and spatially variable environments
- Heterozygosity can increase adaptive plasticity in bacterial cells, in a similar manner to the evolutionary benefits provided by multicopy plasmids.

## Introduction

Bacterial cells typically carry a single (haploid) copy of their genome. While most bacterial species experience transient polyploidy during exponential cell growth^1,2^, several species, including *Buchnera aphidicola*^3^, *Deinococcus radiodurans*^4^, and *Vibrio cholerae*^5^, contain multiple chromosome copies (polyploidy) as a permanent feature of their life-cycle^6,7^. Polyploidy is also observed in several extremophilic archaea^8^ and so-called giant bacteria^9,10^ whose macroscopic cells can contain thousands of chromosomes. In some cases, polyploid chromosomes are identical^3,11^, while in other species the chromosomes are heterogeneous^12^. This gives rise to a form of bacterial heterozygosity that is analogous to the diversity that originates from allelic heterogeneity in different copies of multicopy plasmids^13^. Experiments and theory have shown that plasmid diversity can accelerate adaptation to stress or competition and improve survival in fluctuating environments^14,15^. At the same time, multicopy plasmids are often associated with a metabolic burden for the host cells leading to a fitness cost (plasmid paradox)^16–18^. Despite parallels between multicopy plasmids and chromosomes, the evolutionary consequences of bacterial chromosomal polyploidy and heterozygosity remain poorly understood^19^. Our objective in this study is to address this question using a uniquely tractable experimental system of cell wall-deficient (CWD or L-form) bacteria derived from the actinomycete *Kitasatospora viridifaciens*.

L-forms are bacterial cells lacking the rigid cell wall that maintains cell shape and structure^20,21^. Cell proliferation occurs through asymmetric processes that result from the interplay of membrane synthesis and biophysical forces^22–24^. Hence, populations of L-forms are highly diverse, with cell volumes that can vary by up to two orders of magnitude (Figure 1A). More importantly, because chromosomal segregation is unregulated, cells vary markedly in chromosomal copy number^25^. This mode of replication and proliferation, together with the fact that L-forms can undergo cell-cell fusion^26,27^, results in extensive polyploidy, where multiple chromosomes coexist within the same cell.

**Figure 1:**
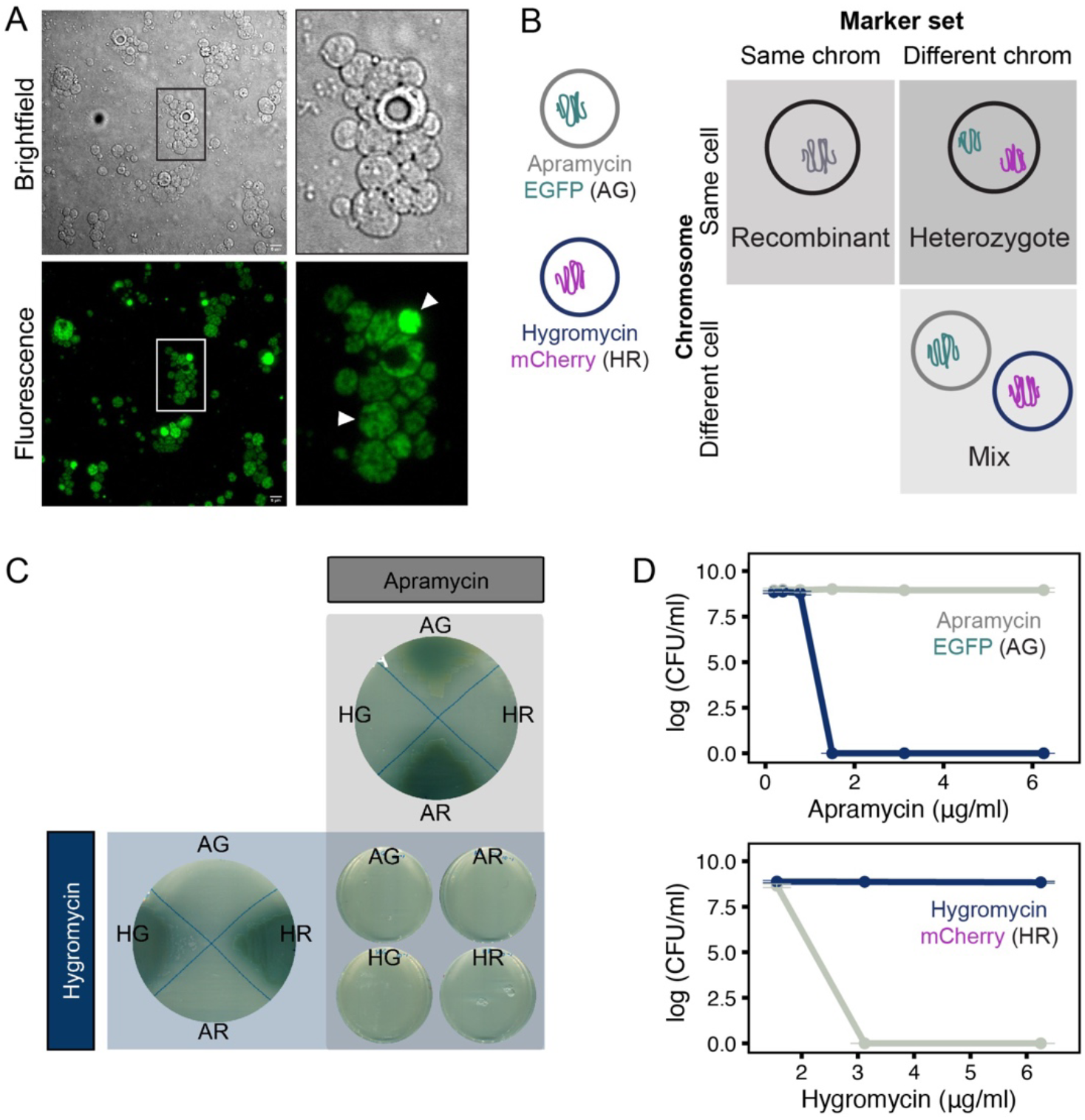
Cell wall-deficient model system. (A) L-forms of wild type *K. viridifaciens* have a spherical morphology like other cell wall-deficient species. Nuclei when stained with SYTO9 (green) show multiple foci in a single cell (white arrow) - a hallmark of polyploidy. (B) Schematic of all strain types generated and used in the study. The strains can be grouped based on the chromosome co-occurrence and marker set (apramycin:EGFP and hygromycin:mCherry). (C) Growth of genetically modified L-forms on selective media (apramycin (A), hygromycin (H), EGFP (G), mCherry (R)). AG and AR show growth (green biomass) only on apramycin while HG and HR show growth only on hygromycin. None of the individual strains can grow in media with dual antibiotic selection. (D) Minimum inhibitory concentration of apramycin (above) and hygromycin (below) for AG (grey) and HR (navy blue) strains, quantified as colony-forming units/ml over a range of tested concentrations.

To examine the effects of chromosomal polyploidy and heterozygosity in L-forms we engineered a pair of chromosomes that could be distinguished by different drug resistance and fluorescent markers^26^. We then generated three combinations (Figure 1B) in which markers were either carried on separate chromosomes in different cells (mixed populations), on a single chromosome (recombinants), or on separate chromosomes within the same cell (heterozygotes). Since all our strain types have multiple copies of each chromosome, we use the terms homozygous to indicate polyploid cells with only one chromosome type (as present in mixed populations and recombinant cells) and heterozygous to refer to polyploid cells with different chromosome types.

We examined growth and survival of these different strain types across a range of antibiotic conditions. Our results provide strong evidence that heterozygous cells persist over a broader range of antibiotic concentrations than homozygotic recombinant cells or mixed populations. Moreover, we found that heterozygous cells can adaptively adjust chromosomal frequencies and allele ratios in response to external conditions. This makes it possible for cells to adapt to antibiotic stress while avoiding costs of expressing both markers from the same chromosome. Our findings indicate that bacterial heterozygosity can benefit cells in fluctuating environments by maintaining genetic diversity, much like multicopy plasmids.

## Results

### Phenotypic characterization of wall-deficient cells

Cell wall-deficient L-forms of *K. viridifaciens* (Figure 1A) were genetically modified to carry combinations of an antibiotic resistance gene (either conferring resistance to apramycin (A) or hygromycin (H)) and a fluorescence marker gene (either (EGFP(G) or mCherry(R)). This resulted in four strains: AG, AR, HG, and HR (as illustrated in Figure 1B, 1C). Resistance phenotypes were as expected, with high-level resistance to each respective resistance allele and no cross-resistance (Figure 1C). Apramycin resistant strains had an MIC of 3.1 µg/ml for hygromycin while hygromycin resistant strains had an MIC of 1.6 µg/ml for apramycin (Figure 1D). Neither strain grew on media with both antibiotics (Figure 1C). The AG and HR strains were subsequently fused to create a heterozygous (H) strain containing both chromosomal types within a single cell (Figure 1B). On the other hand, wildtype strain was consecutively transformed to generate a recombinant (R) strain containing all four markers on a single chromosome (Figure 1B).

Growth and fluorescence of the heterozygous (H), recombinant (R) or mixed (M) (which refers to a 1:1 mixture of AG and HR monocultures) populations was assessed by spotting 3 μl from a 0.1 OD_600_ culture in different environments varying in the presence or absence of apramycin or hygromycin (Figure 2). In the absence of antibiotics, mixed populations showed patches of either fluorescence colour (Figure 2A), with some colocalization of fluorescence across the spot; colocalization was quantified as the correlation between the pixel distribution across the two channels (EGFP and mCherry) (Figure 2, bottom panels). Growth on either antibiotic alone led to monomorphic populations expressing only a single fluorescence marker, indicating that fluorescence is a good proxy for the linked resistance allele. Neither strain in the mixed combination persisted when grown in the presence of inhibitory concentrations of both antibiotics.

**Figure 2:**
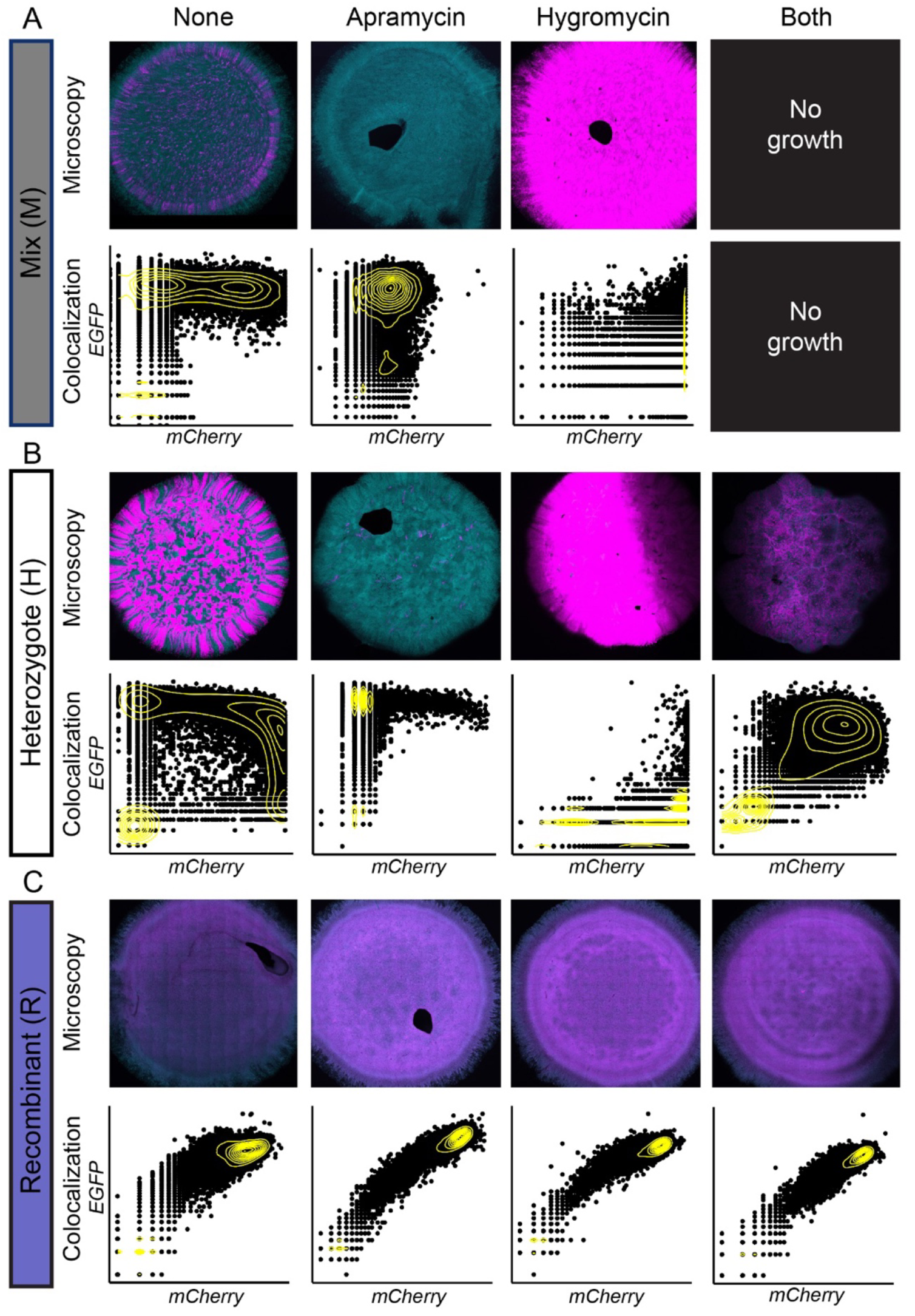
Fluorescence marker expression of strains across antibiotic selection. Different strain types (A) Mixed, (B) Heterozygote and (C) Recombinant grown with or without both antibiotics, only apramycin, or only hygromycin. Fluorescence microscopy of individual colonies (top rows) and the corresponding quantification of fluorescence (bottom rows) shows the expression and colocalization of EGFP (cyan) and mCherry (magenta) in different strains in response to selection. Pixel-wise colocalization of either marker was quantified for each pixel within the colony area and 10,000 points chosen at random were plotted. A scatter pattern tending towards either the x (mCherry) or y (EGFP) axis indicates dominance of one marker, whereas a distribution in the top right region indicates colocalization of cells expressing both markers.

In contrast, heterozygous and recombinant cells grew under all antibiotic exposures. In the absence of selection, heterozygous populations showed patches/sectors of both EGFP and mCherry, suggesting that either chromosomal loss results in homozygotic lineages, or there is a shift in the ratio of chromosomes containing different markers. We consider the latter possibility in the next section. In the presence of both antibiotics (Figure 2B, 2C, last column), we observed significant colocalization of EGFP and mCherry signals across the colony, consistent with the maintenance of both chromosome types (Pearson’s R_heterozygote_ = 0.487, R_recombinant_ = 0.885 Figure 2B bottom panel). During single antibiotic selection, heterozygous populations, like the mixed populations, shifted towards the dominance of a single fluorophore, suggesting that the unselected marker was lost or reduced in frequency (Pearson’s R_heterozygous_ = 0.057 (apramycin), R_heterozygous_ = 0.28 (hygromycin)). Recombinant cells grew in all conditions, with significantly more colocalization of the fluorescent markers than heterozygous cells (Pearson’s correlation coefficient ranging from 0.663 to 0.917, figure 2C bottom panel). These results confirm that the strains respond predictably to selection for growth under antibiotic exposure and in terms of fluorescent marker expression.

### Heterozygous cells persist across a broader range of antibiotic concentrations

To measure the effects of heterozygosity in more detail we examined cell growth across a gradient of antibiotic concentrations from 0 - 2x MIC, as shown in the schematic (Figure 3A). The resulting colonies were assessed by quantifying CFUs (Figure 3B, 3C, 3D) and with microscopy (Figure 3E, 3F). At sub-MIC concentrations in mixed populations, fluorescence intensity of EGFP and mCherry showed marked deviations, consistent with distinct populations resulting from the loss of the linked drug resistance genes (Figure 3E). We observed only sparse growth at and above the MIC (Figure 3E, dotted box, Supplementary Figure 1A).

**Figure 3:**
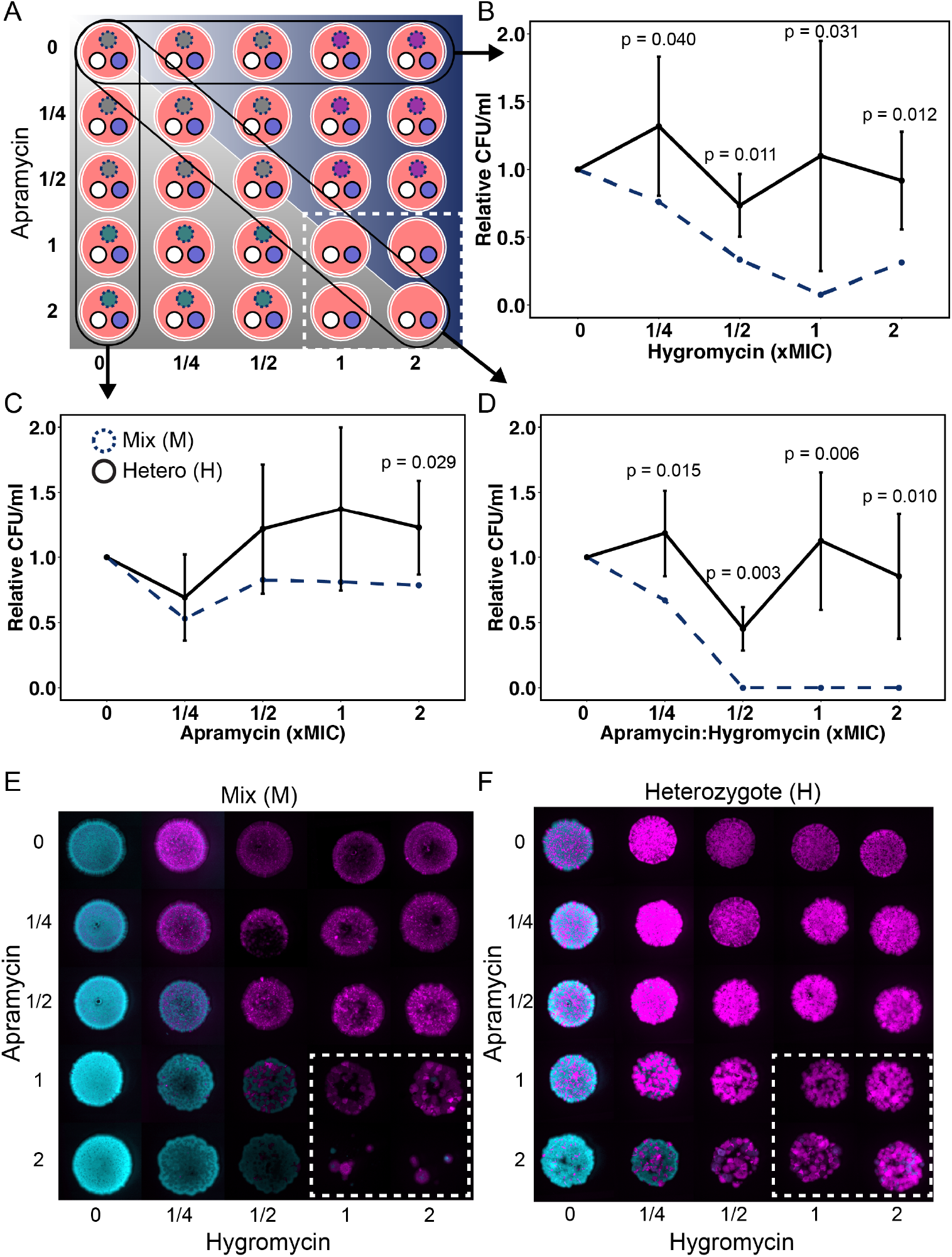
Higher growth range due to heterozygosity. (A) Experimental set-up of individual plates consisting of varying concentrations of apramycin and hygromycin below and above MIC (0x-2x). Each plate was inoculated with three strains (Mixed (dotted circle), Heterozygote (smooth white circle), and Recombinant (smooth purple circle)). Each spot represents the predicted growth (presence or absence) and fluorescence expression (cyan for EGFP and magenta for mCherry). (B, C, D) After 3 days of growth, appropriate dilutions were used to quantify cell numbers by plating. Heterozygotes (bold lines) show growth at all concentrations while Mixed populations do not (dotted lines). Significant differences are reported for each tested pair of Heterozygote and Mixed populations (Welch t-test, n=3). (E, F) Fluorescence images of colonies after 3 days of incubation. (E) The Mixed (M) strain grows well below MIC with either EGFP or mCherry expression corresponding to the antibiotic selection. Growth is significantly reduced or absent when exposed to both antibiotics above the MIC (dotted white box). (F) The Heterozygote (H) strain can grow equally well across the gradient below and above MIC. Markers not under selection are visible in patches, consistent with shifts in chromosome/allele frequencies.

By contrast, heterozygous cells grew well in all antibiotic concentrations (Figure 3B, 3C, 3D). The benefit of chromosomal diversity in heterozygous cells is shown in two ways. First, we observed a significant relative advantage of heterozygous versus mixed cells in terms of CFU (Figure 3B, 3C, 3D, Welch t-test). Second, in contrast to mixed populations, heterozygous cells grew well in the presence of both antibiotics at concentrations at and above the MIC, as indicated by correlated patches of both fluorescent markers (Figure 3F, Supplementary Figure 1B). Interestingly, the advantage to heterozygous cells was observed both above and below the MIC, for single and double antibiotic exposure. This indicates that heterozygous cells grow better over a broader environmental range than either genotype in the mixed populations.

Recombinant colonies show growth all across the antibiotic gradient with an even expression of both fluorescence markers (Supplementary Figure 1C). However, in comparison to recombinant cells, the growth of heterozygous cells in colonies is less spatially uniform suggesting the possible loss or skew in frequencies of either chromosome type (Supplementary Figure 2).

### Fixed versus flexible costs of antibiotic resistance

If recombinant populations grow across the same range of drug concentrations as heterozygous cells, what prevents their fixation? One factor is the potential cost of constitutively expressing both drug resistance markers, especially when they are not under antibiotic selection^16^. To quantify these costs, we estimated the fitness of the recombinant strain when competing against either parental strain: AG (Figure 4A) or HR (Figure 4B). Regardless of the initial frequencies (10:90, 50:50, or 90:10), we observed that the recombinant strain was rapidly outcompeted by the parent strains (Figure 4C), consistent with a significant metabolic cost to antibiotic resistance.

**Figure 4:**
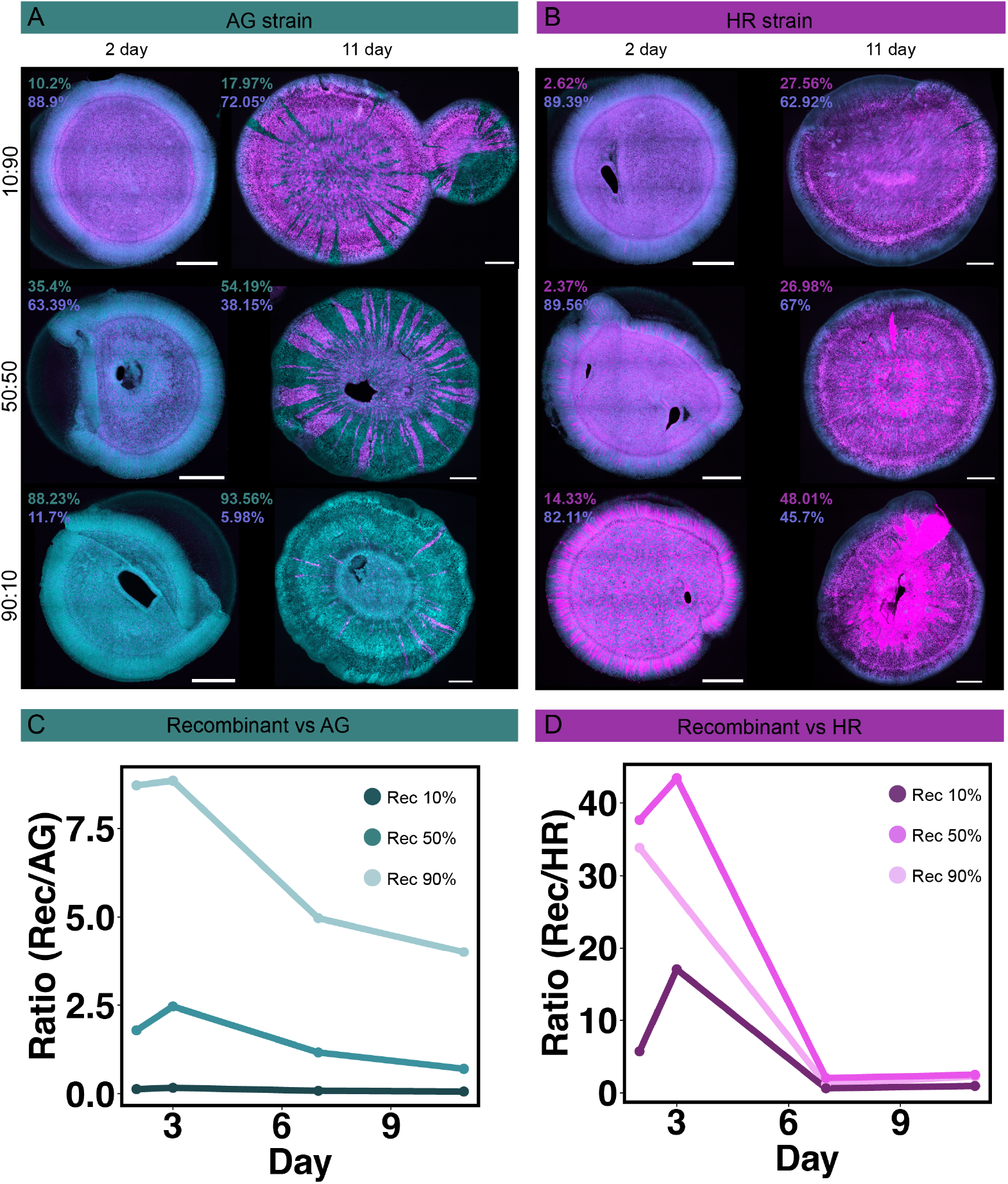
Fitness cost for recombinant cells. Parental strains (A) AG or (B) HR were mixed with the Recombinant (R) in different starting ratios (10:90, 50:50, 90:10) and incubated for 11 days. Fluorescence microscopy of individual colonies at different time points (representative images from days 2 and 11 are shown) were used to quantify the ratio of EGFP (cyan) and mCherry (magenta) expressing cells indicated as percentages next to the colony image. Based on this measurement, the ratio of (C) R to AG or (D) R to HR was calculated over time, with the Recombinant declining at all inoculation frequencies.

In contrast to the fitness costs observed in recombinant strains, we hypothesised that these costs could be reduced in heterozygotes by flexibly altering chromosome (resistance allele) ratios in response to external conditions. To test this, we used quantitative PCR to quantify the frequencies of antibiotic resistance markers (*aac(3)IV* and *hph* for the apramycin and hygromycin resistance genes, respectively) in mixed, heterozygous and recombinant cells grown across a gradient of both antibiotics.

Our results show how the frequency of each antibiotic resistance gene changes in the three cell types after exposure to different antibiotic concentrations (Figure 5A, Supplementary Figure 3). These results lead to the following conclusions. First, antibiotic resistance gene frequencies are responsive to environmental conditions (Figure 5, Supplementary Figure 3A, B, C). At the highest concentrations of both antibiotics, mixed and heterozygous populations showed strongly skewed frequencies of each resistance allele. In the mixed population this skew was due to selection of either resistant strain by the corresponding antibiotic and the extinction of the other (Supplementary Figure 3B). By contrast, in the heterozygous cells, selection acted between chromosomes within each cell, resulting in a skewed chromosomal ratio at the per cell level that corresponded predictably to the associated antibiotic exposure (Supplementary Figure 3C). Importantly, while this shift caused a significant loss of viability in the mixed population (Figure 3), heterozygous cells retained normal growth. Second, while resistance markers in mixed populations were entirely lost during exposure to the highest concentration of either antibiotic alone (ranging from 0-100%), they were maintained in heterozygous cells across all drug exposure conditions (ranging from ∼3-97%) (Figure 5B). Thus, while growth of mixed populations was curtailed due to the loss of functional diversity, the maintenance of both alleles in heterozygous cells permitted their growth across a broader environmental range. By contrast, resistance gene frequencies in recombinant cells were essentially invariant, regardless of conditions and associated costs (Supplementary Figure 3D).

**Figure 5:**
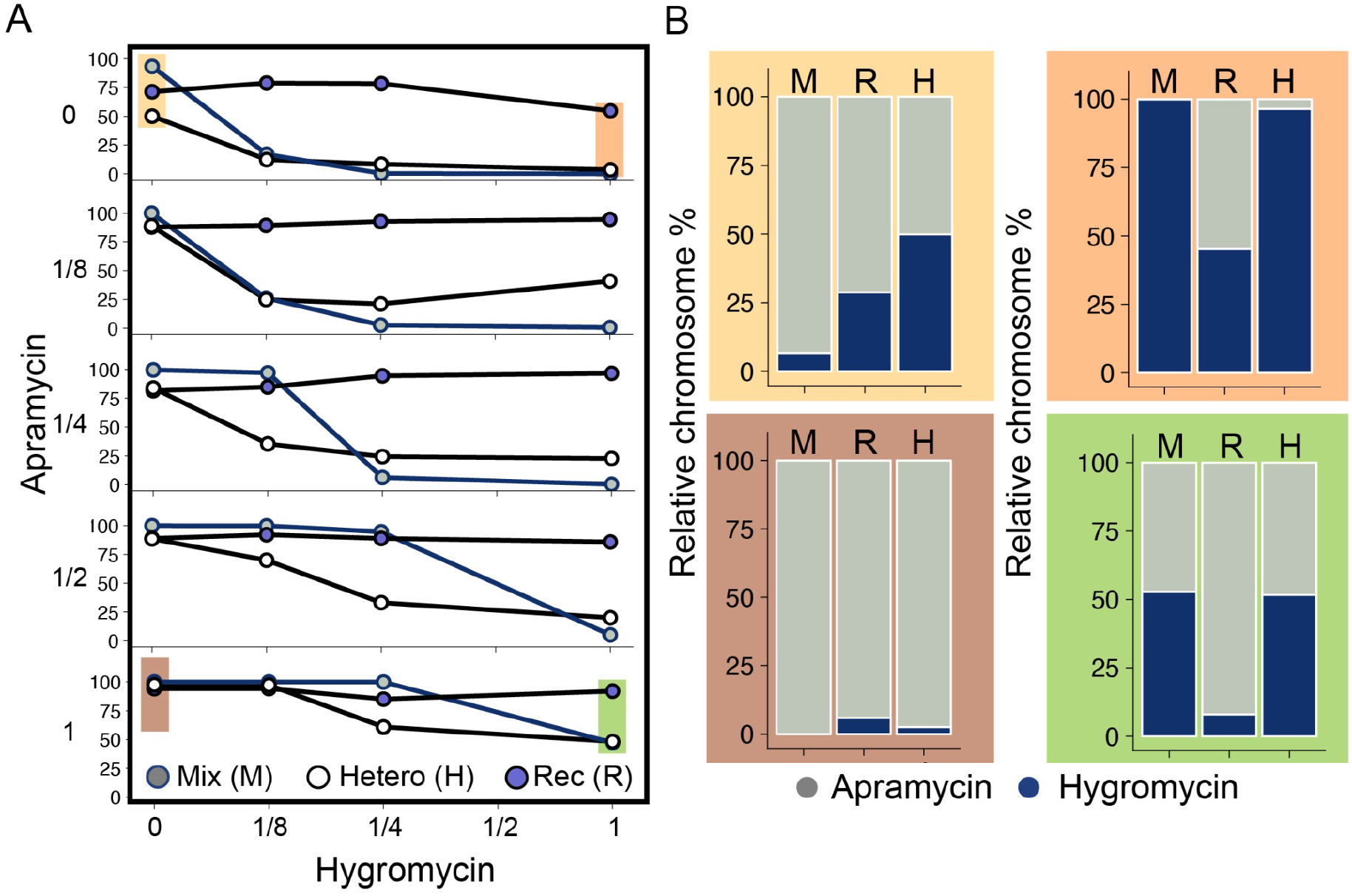
Marker ratio change over the selection gradient. (A) The frequency of apramycin and hygromycin resistance markers was quantified via qPCR from the collected biomass of Mixed (M), Recombinant (R), and Heterozygote (H) populations. The ratio of apramycin to hygromycin marker shows different patterns across the gradient of selection for the Mixed (grey) and Heterozygote (white) populations, while the ratio in Recombinant (purple) populations is consistent across antibiotic exposure levels. (B) The relative chromosome percent at the extreme antibiotic concentrations shows the maintenance of both markers in the Heterozygote population even at extreme concentrations of either antibiotic alone. In the highest hygromycin region (orange box), the apramycin marker (grey bar) is maintained, and in the apramycin high region (brown box), the hygromycin marker (navy blue bar) is maintained. This suggests that the Heterozygote population is able to retain both resistance markers even under extreme selective pressure.

### Heterozygotes show segregated response to temporal and spatial antibiotic gradients

Resources and stress vary spatially and temporally in natural environments. We next asked if heterozygous cells could flexibly adjust chromosome ratios to persist in these dynamic environments. To address this, we examined cell growth and fluorescence expression (indicative of allele frequencies) on plates with opposing antibiotic gradients (Figure 6A and Supplementary figure 4A). Because diffusion from paper disks containing antibiotics occurs continuously over time, this approach allowed us to analyse the effects of heterozygosity as the colony grew.

**Figure 6:**
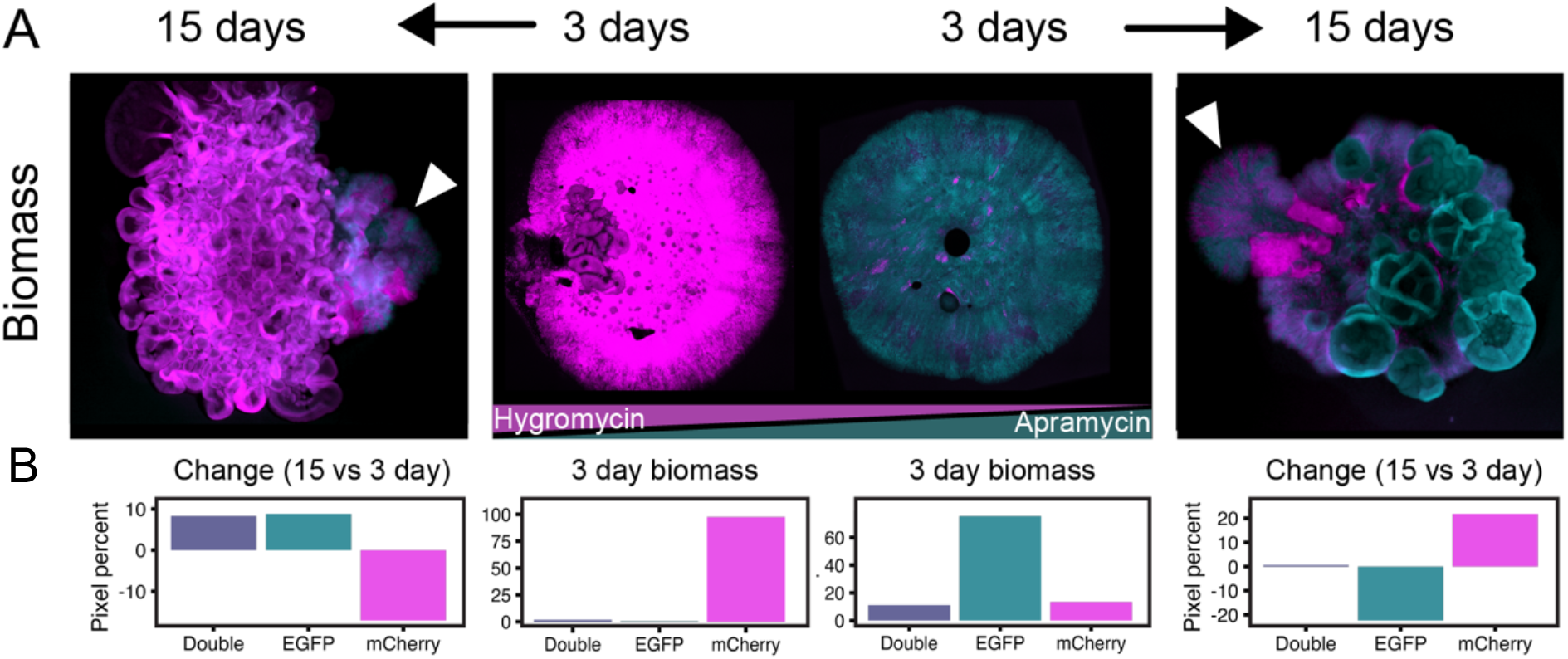
Heterozygous population shows a spatio-temporal response to opposing antibiotic selection gradients. (A) The heterozygous population was exposed to an opposing gradient of hygromycin and apramycin. (B) Fluorescence markers were used to visualize and quantify the population’s response, with mCherry (magenta) used as a marker near the hygromycin gradient and EGFP (cyan) used as a marker near the apramycin gradient. At 3 days, the fluorescence pixel percent correlated with the antibiotic selection, with high mCherry (97%) near the hygromycin gradient and high EGFP (75%) near the apramycin gradient. From 3 to 15 days, the fluorescence pixel percent showed an opposite trend, with an increase in EGFP (8.8%) near the hygromycin gradient and an increase in mCherry (21.8%) near the apramycin gradient.

Three days after cells were inoculated onto plates with opposing antibiotic gradients, heterozygous cells experienced a high concentration of one antibiotic and a low concentration of the other. As expected, fluorescence intensities corresponded to the level of drug exposure (Figure 6A): cells grown near apramycin (shown in green) predominantly expressed EGFP (75%) while cells grown near hygromycin (shown in pink) overwhelmingly expressed mCherry (97%) (Figure 6B). However, after 15 days, during which time the concentration of both antibiotics changed due to diffusion (Supplementary Figure 4A), we observed a corresponding shift in fluorescence expression. As the concentration of hygromycin declined, so did the expression of mCherry (from 97% to 80%). At the same time, because cells were increasingly exposed to apramycin, there was a corresponding increase in the expression of EGFP (from 0.6% to ∼ 9%). The same qualitative response was seen in cells initially grown closer to apramycin, although to a lesser degree due to the different concentration-dependent behaviour of this antibiotic (Figure 1). As the concentration of apramycin decreased, the expression of EGFP also decreased (by 22.4%) while as hygromycin selection increased the expression of mCherry increased (by 21.8%). For both cases (biomass initially close to hygromycin or close to apramycin), the fraction of double-labelled cells increased most visibly on the side of the colony exposed to both antibiotics (white arrows).

Parallel experiments in antibiotic gradients were carried out with mixed and recombinant populations. As expected, mixed populations were growth arrested over 15 days due to exposure to both drugs (Supplementary Figure 4B); while recombinant cells grew but their marker expression was invariant (Supplementary Figure 4C).

## Discussion

Bacterial polyploidy creates opportunities to increase genetic diversity within a single cell by retaining chromosomal copies^12,13,28,29^. To understand the evolutionary consequences of this phenomenon, we used cell-cell fusion in *K. viridifaciens* L-forms to create heterozygous lineages that were then examined across fixed and dynamic gradients of antibiotics. Our results show that heterozygosity allows cells to persist across a broader range of antibiotic concentrations than homozygous cells (recombinant or a mixed population). In addition, heterozygotes can offset costs of antibiotic resistance by flexibly adjusting the frequency of resistance genes to match external drug exposure. This response occurs rapidly in the presence of a static antibiotic concentration and dynamically in response to a shifting antibiotic gradient. Together, our results suggest that the plasticity provided by chromosome heterozygosity may facilitate bacterial adaptation to environmental heterogeneity or stress.

The benefits of bacterial heterozygosity mirror effects of analogous systems in bacteria that increase genetic diversity^2,9^, gene expression^30^ or protein abundance^31^ through increased copy number. For individual genes or smaller chromosomal regions, tandem gene amplifications can lead to antibiotic heteroresistance^2,14,32^; this allows otherwise susceptible cells to become transiently drug tolerant while also increasing the number of mutational targets that can lead to more stable resistance. Multicopy plasmids offer similar benefits in high antibiotic concentrations or across fluctuating environments with varying levels of antibiotic exposure^13^.

In each of these analogous systems, the benefits afforded by increased copy number or gene expression allows flexible and plastic responses whose effects are more rapid than genetic changes via mutation. At the same time, the costs of overexpression or plasmid carriage ensure that these processes are transient and highly variable across bacterial populations. Heteroresistance is only observed in a minor fraction of bacterial cells, while plasmid copy number can vary markedly depending on the concentration or frequency of antibiotic exposure^14,32^. Similarly, our results suggest that the benefits of chromosome heterozygosity are not consistent across all conditions and may also be transient. During antibiotic exposure, chromosome copy number can rapidly shift to increase the frequency of the appropriate resistance gene. However, just as tandem duplications and plasmids are lost, so too are chromosomes due to unequal segregation during cell division. As far as we are aware, there are no mechanisms to ensure reliable chromosome partitioning or copy number in L-form cells. In L-forms, segregational loss can potentially be overcome by cell-cell fusion that would recreate a new heterozygous lineage^26,27,33,34^. This type of fusion and segregation is thought to have been a major driver of adaptability of protocells that existed before the evolution of the more rigid bacterial cell wall^21^.

Numerous processes have evolved to provide microbial populations with the ability to generate adaptive plasticity in response to environmental stress or instability. Our results suggest that chromosomal heterozygosity offers bacteria yet another mechanism to flexibly respond to shifting environments. Further work will be needed to assess how widespread this phenomenon is across microbial groups and environments.

## Materials & methods

### Strains and media

Apramycin resistant:GFP (AG) and hygromycin resistant:mCherry (HR) lines of *K. viridifaciens* DSM40239 were used from a previous study which details the strain construction^21^. Briefly, plasmids containing the marker combinations along with a ϕC31 *attP* site and integrase gene were introduced into L-forms via PEG-induced transformation. The resultant transformants were directly selected on antibiotic containing media. For the fusion strain, a poly(ethylene) glycol (10%) mediated fusion was used to fuse AG with HR followed by two rounds of selection on both antibiotics. The resultant strain contains different chromosome types in the same cell with the marker sets on separate chromosomes. The recombinant strain was obtained by transforming a wildtype *K. viridifaciens* L-form with both sets of markers in consecutive steps such that insertion takes place on the same chromosome (Figure 1B). Transformation was carried out using a modified protoplast transformation protocol using plasmids containing the two marker sets. The first insertion takes place at the *attP* site (via ϕC31 integrase) and the second insertion potentially occurs at a *pseudo-attP* site^35^. This recombinant strain contains all marker sets on the same chromosome and was enriched by selection on plates containing both antibiotics at least 3 times. All strains were grown on L-phase broth (LPB) which consists of a 1:1 mixture of yeast extract malt extract and tryptic soy broth supplemented with 10% sucrose and 25 mM MgCl_2_. Colony growth for quantification and imaging was done on L-phase agar (LPMA) which is LPB containing 1.5% agar, 5% horse serum and 25 mM MgCl_2_. Apramycin and hygromycin (Duchefa Biochemie) were added as per experimental requirement and all incubations were carried out at 30°C.

### Minimum inhibitory concentration (MIC) quantification

Monocultures of AG and HR were grown in LPB for 3 days from a stock culture. These cultures were adjusted to 0.1 OD_600_ and spot diluted (1:10 over 6 rounds) on LPMA plates containing increasing amounts of antibiotics. For apramycin the range tested was (in decreasing order): 6.25, 3.12, 1.56, 0.78, 0.39, 0.19 μg/ml. For hygromycin the range tested was (in decreasing order): 50, 25, 12.5, 6.25, 3.12 and 1.56 μg/ml. Both strains were plated on all concentrations and incubated for 3 days. Colonies in spots were counted and CFU/ml was quantified.

### Competition experiment

Pairwise mixes of OD_600_ adjusted cultures were prepared with varying ratios of parent (AG or HR) to recombinant strains (10:90, 50:50 and 90:10), to control for frequency-dependent effects. Each mix was spotted onto media without antibiotic selection to test for the cost of carrying resistance markers. The spot was allowed to grow over 11 days and imaging was done at day 2 (colonies become visible), 3, 7 and 11. Images were processed and analysed for percentage of pixels in both channels (EGFP and mCherry) to track the parental population over time. The single channel pixel percent (EGFP or mCherry) tracks the parental strain ratio while the overlapping pixel percent (EGFP and mCherry) is representative of the recombinant strain.

### Cell growth across a range of antibiotic concentrations

Population growth (fused and mixed) was tested across a range of antibiotic concentrations (0, ¼, ½, 1 and 2x MIC of apramycin and hygromycin). Cell culture of 3 μl in the form of a spot was grown at each specified antibiotic concentration in a plate consisting of defined concentrations of apramycin and hygromycin as shown in figure 3A. Each condition and strain were tested in triplicate. After 3 days of growth at 30°C we quantified CFU/ml using the appropriate dilution and isolated single colonies for microscopic analysis.

### Microscopy and image analysis

Whole colony imaging was done by cutting out and flipping the colony into an Ibidi® 8-well chamber slide. An imaging set-up of a Zeiss LSM 900 Airyscan 2 microscope was employed to visualize the fluorescently labelled microbial strains at a magnification of 40x. EGFP strains (AG, Rec, Fused) were excited at a wavelength of 488 nm, with emission detection at 535 nm. Conversely, the mCherry strains (HR, Rec, Fused) were excited at 535 nm, and its emission was captured at 650 nm. Utilizing the Zen software developed by Zeiss, we acquired multichannel images encompassing both fluorescence and bright-field signals in a multistack format. These images were subsequently subjected to analysis using ImageJ/Fiji^36^.

Multiple tiles were imaged to cover the entire microbial colony. These tile images were then stitched together with 10% overlap. Within each fluorescence channel, a thresholding process was applied to delineate the overall pixel area. These thresholded images were subsequently utilized for two distinct calculations: (i) determination of the total area, accomplished by applying the OR function in the Image calculator. (ii) Computation of the fused area, achieved through the application of the AND function. The total area selection enabled us to quantify the individual pixel areas occupied by both green and red pixels, as well as those occupied by either green or red pixels. Using the Coloc2 plugin, a correlation was calculated and plotted to identify the trend in pixel intensity between the green and red channels for different images.

For greater resolution of strains within colonies a confocal microscope was used. The colony along with solid media was cut out from the plate and flipped over onto an ibidi® slide ensuring no air bubbles were trapped between colony biomass and the slide. Imaging was performed from an inverted objective with multiple z-axis to cover the outer edges as well as the bulging center. Images were taken with the fluorescence setting for EGFP and mCherry.

### Quantitative PCR based chromosome ratio estimation

Biomass of a single spot was collected and resuspended in lysis buffer. The microfuge tubes were then heat treated (90°C for 10 minutes) to break open cells for DNA extraction. The pellet was centrifuged and supernatant diluted (1:10) for qPCR analysis. RT-qPCR was done using the iTaq Universal SYBRGreen Supermix kit (Biorad) following the vendor protocol. The reaction mix consisted of 2 μl DNA (supernatant from above), 1 μl forward primer, 1 μl reverse primer, 10 μl qPCR mastermix (contains dNTPs, enzyme and buffer), 1 μl DMSO (used for high GC DNA in Streptomyces) and 5 μl water. The primers (Supplementary Table 1) were used at a working concentration of 0.5 uM. The target was approximately 150 bp and a range of 1 ng to 0.1 pg DNA mass was used for the standard curves. The following thermocycler program was used: 95°C for 5 min, 39 cycles of [95°C for 5s, 60°C for 30s, and 65°C for 5s], followed by 95°C for 30s.

Purified amplicons of the apramycin resistance gene (*aac(3)IV*) and hygromycin resistance gene (*hph*) were used for generating standard curves, *infB* was used as a housekeeping gene for reference. Standard curves were generated (Supplementary figure 5) for the house keeping gene (*infB*), apramycin resistance gene (*aac(3)IV*) and hygromycin resistance gene (*hph*). A serial 10-fold dilution was carried out of the pcr product followed by qPCR and plotting ct values obtained for the known copy number (Supplementary Figure 5). R^2^ values indicated that the Ct values can be used to accurately predict the copy number of the corresponding gene given the same PCR settings. The above slope values were used to calculate the relative proportion of apramycin and hygromycin resistance genes in samples across the antibiotic gradient. In each run purified pcr products (mentioned above) were used as reference and the sample (DNA obtained from the biomass) was the test. The following formulas were used-

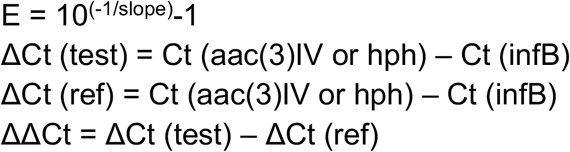

Relative chromosome number (RCN) = (1 + E)^-ΔΔCt^

### Resistance and growth in antibiotic gradients

To create an antibiotic concentration gradient on the plate, filter discs were prepared with the corresponding antibiotic concentrations (50 μg/ml apramycin, 100 μg/ml hygromycin). These filter discs were then placed 1 cm from the bacterial inoculum. The layout of filter discs with different antibiotics around the inoculated spot is shown in Supplementary Figure 5A. The plate was then kept in the incubator for 1 hour followed by inoculation of different strain types. After inoculation colony growth and fluorescence were tracked over time (3 to 15 days).

### Statistical analysis

All statistical analysis was performed using R Statistical Software (v4.2.1; R Core Team 2022). Significant differences and tests used have been mentioned in the results where applicable. Comparison of Mixed, Recombinant and Heterozygous strains when done in pairs was by using the Welch t test. For image analysis, pixel-wise correlation significance was calculated using Pearson’s coefficient.

## Acknowledgments

We thank members of the Rozen lab and Claessen lab for fruitful discussions and suggestions. SS acknowledges Samir Giri and Jordi van Gestel for comments on the manuscript. SS was supported by the Origins Centre (NWA impulse) for funding. TvD acknowledges NWO for funding. DC and DR were supported by the Dutch Science Foundation (NWO: ALW).

## Author contributions

SS, DC and DR conceived and designed the study. SS and TvD performed the experiments and analysed the data. SS wrote the first draft of the manuscript. All authors revised and contributed to the final version of the manuscript.

