## Supplementary figures 1-5, table 1 for "Bacterial heterozygosity promotes survival under multidrug selection"

**Figures- 5 (pages 2-6)**

**Tables- 1 (page 6)**

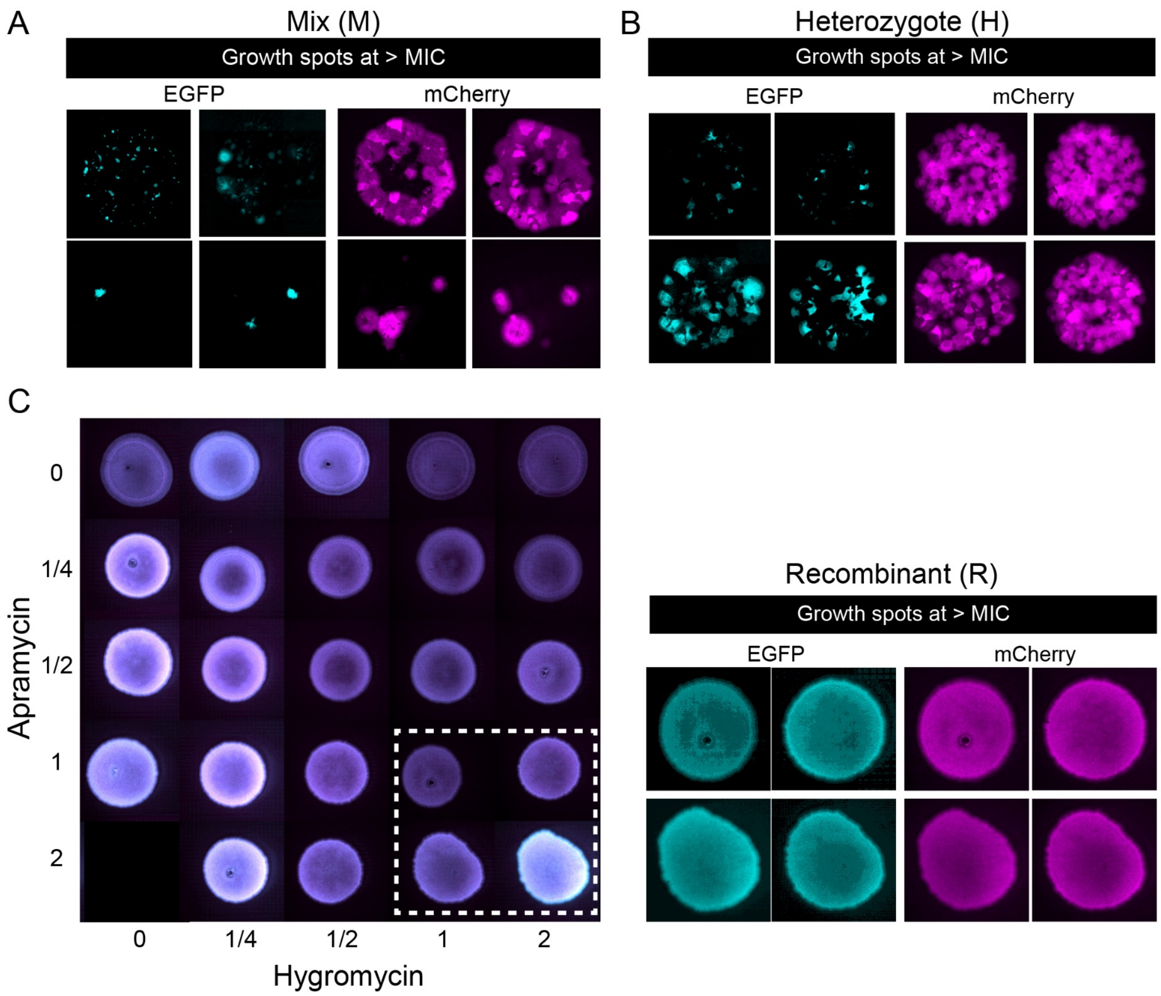


**Supplementary figure 1:** (A, B) Individual fluorescence channel images of the colonies above MIC (dotted white box from Figures 3E and F) showing higher growth and balanced expression in Heterozygotes compared to the Mixed population. (C) Fluorescence images of the Recombinant strain growing across the antibiotic gradient matrix (as shown in Figure 3A). The expression of EGFP and mCherry is uniform across the gradient, which is also evident in the individual channel images of a select few colonies (dotted white box).

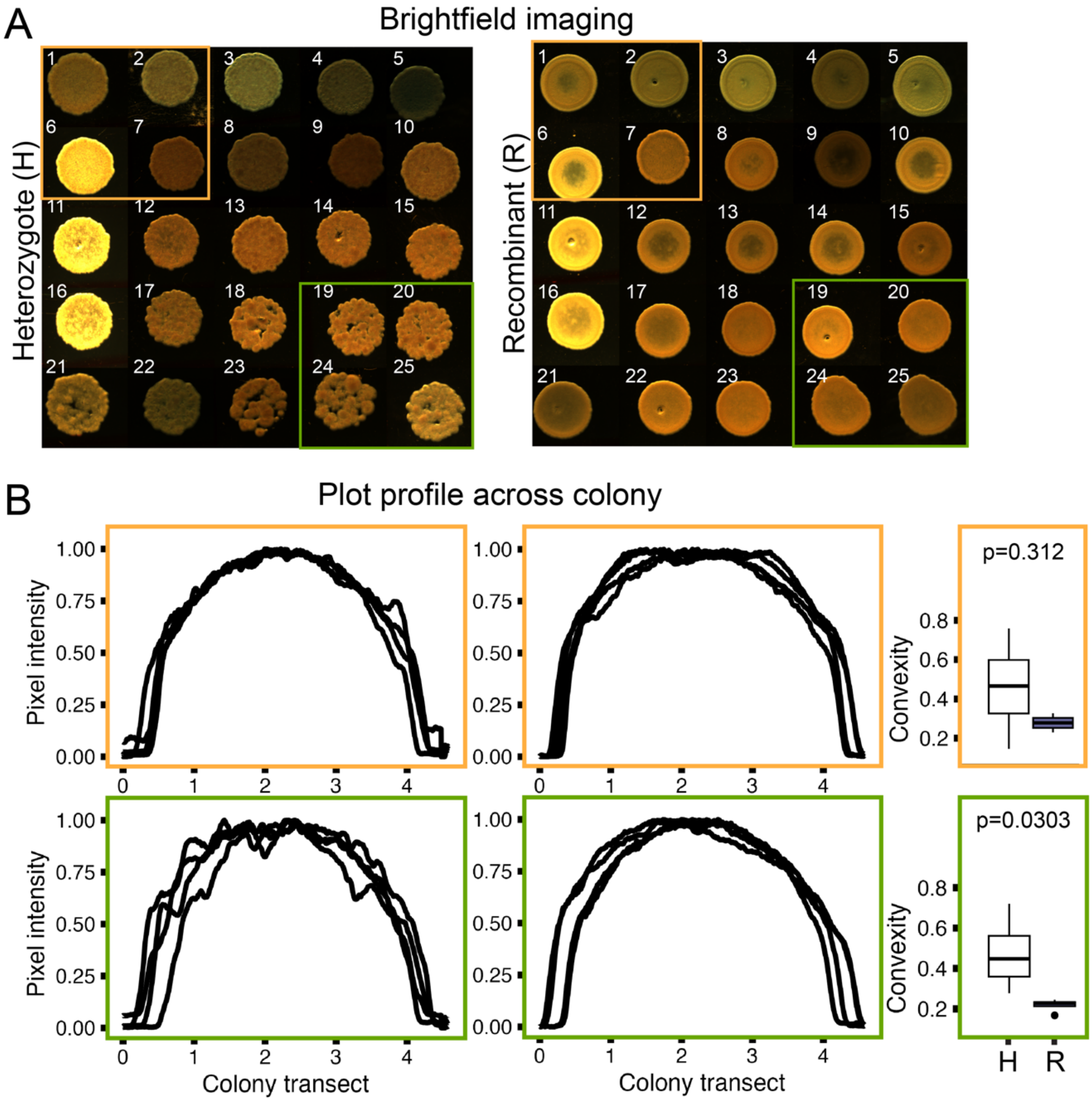


**Supplementary figure 2: Differences in surface architecture of strains**. (A) Brightfield images of colonies taken from the experiment in figure 3. (B) These colonies were analyzed for ruggedness via surface profile plots across the colonies below MIC (orange box) and above MIC (green box). Heterozygote colonies (left panel) show patchiness/ruggedness which is more visible above MIC (green box), while Recombinant colonies (right panel) show consistent, smooth texture across the gradient (convexity score closer to 0). Convextity of all 4 colonies has been plotted for comparison with significant difference above MIC (Welch t test).

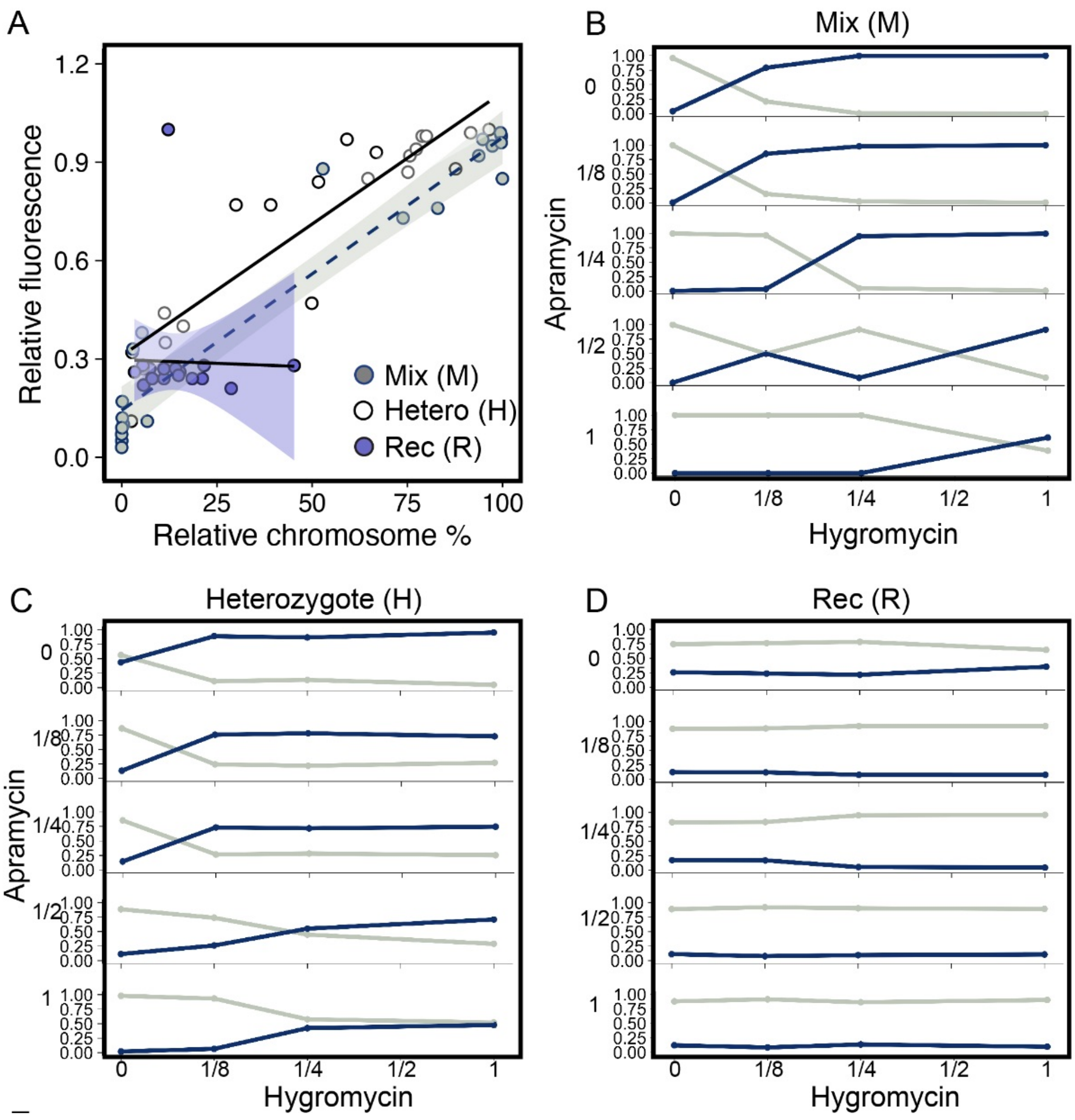


**Supplementary figure 3: Correlation between relative chromosome percent and relative fluorescence expression**. (A) A positive trend was observed between chromosome percent and fluorescence expression for Mixed (grey) and Heterozygote (white), indicating that fluorescence images corroborate well with the chromosome ratios quantified using qPCR. The Recombinant strain (purple) shows a flat trend corresponding to the constant ratio of AG:HR and equal expression of EGFP:mCherry across the tested gradient (figure 3A). (B, C, D) Chromosome percent of AG (grey) and HR (blue) for each strain type across the tested gradient. These values were used to calculate the chromosome ratio that is shown in Figure 5A.

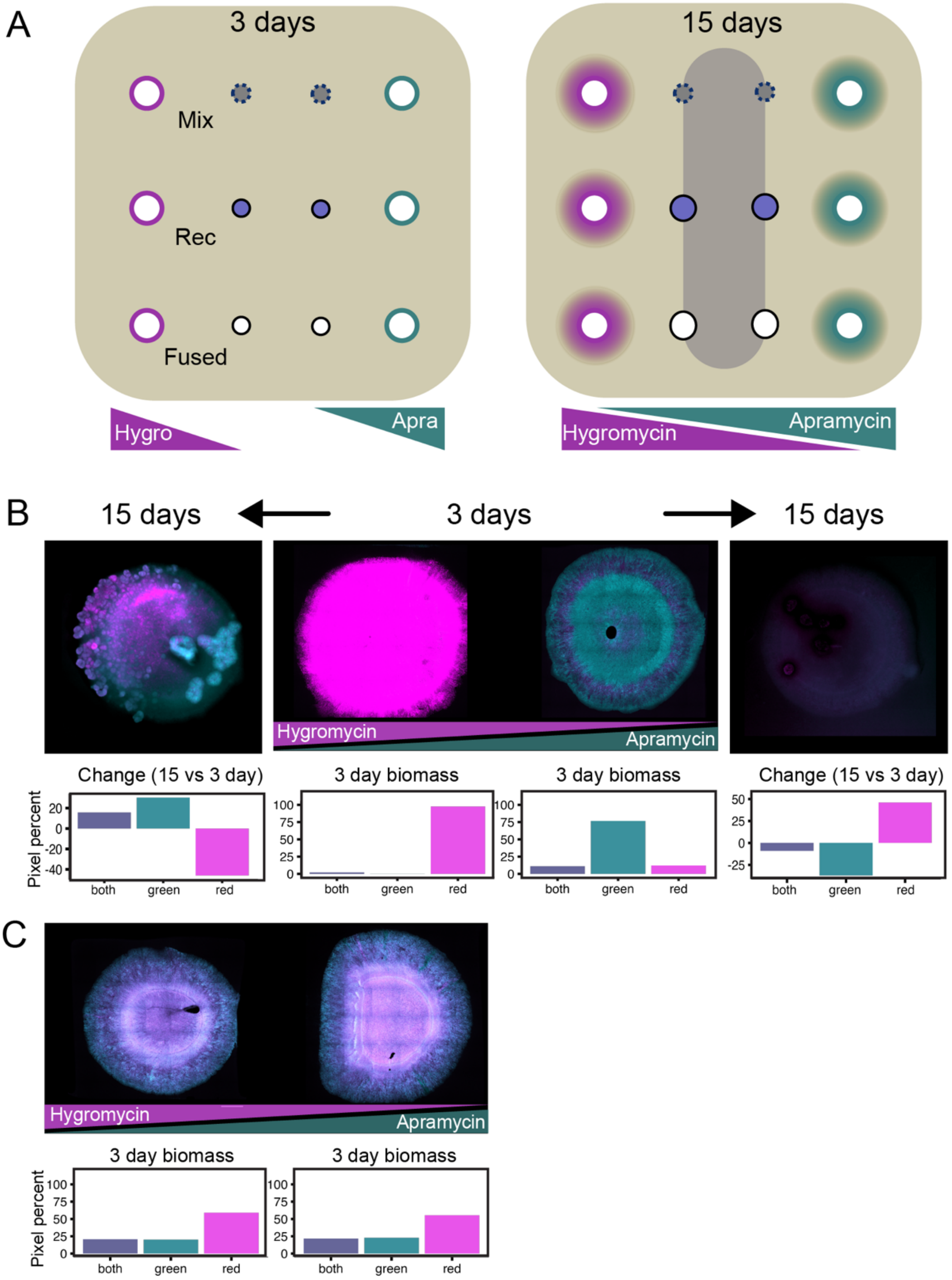


**Supplementary figure 4:** (A) The antibiotic gradient was set up in the same environment by placing filter discs containing antibiotics such that they diffused into the agar medium, indicated by a halo. Populations (Mixed, Heterozygote, and Recombinant) were inoculated in the center, thus experiencing an increasing gradient of a different antibiotic on each side and, over time, also a predcted zone with both antibiotics (purple shaded area). (B) Fluorescence images of colonies of the Mixed populations over time in response to an opposing gradient of hygromycin-apramycin. The fluorescence images were used to calculate the percent of marker expression in each biomass, as shown in the bar plots below. (C) Recombinant population exposed to an opposing hygromycin-apramycin gradient at day 3 do not show any phenotypic difference in marker expression.

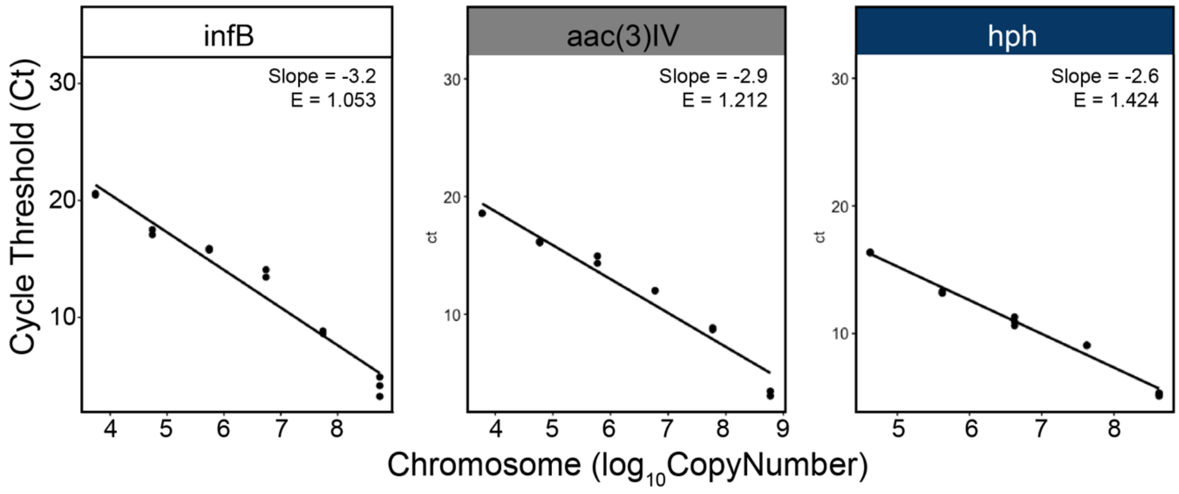


**Supplementary figure 5**: Standard curves for Ct value correlated to chromosome copy number. *infB* gene was used as a reference, *aac(3)IV* gene was used for apramycin and *hph* gene was used for hygromycin.

### Supplementary table 1. Primer sequences used in this study

| **Name** | **Sequence 5'-3'** |
|---|---|
| infB_forward | CGAAACGCCAGGAATATGAAG |
| infB_reverse | CGCCCAGGTTAAACATCAC |
| Apra_forward | GCAATACGAATGGCGAAAAG |
| Apra_reverse | AGATGATCTGCTCTGCCTG |
| Hyg_forward | ACCGTGCTCACCCCCCATTC |
| Hyg_reverse | CCGGAAGGCGTTGAGATGCAG |
